# Cigarette smoke and biological age induce degenerative heterogeneity in retinal pigment epithelium

**DOI:** 10.1101/2025.01.27.635096

**Authors:** Krishna Singh, Yang Jin, Ming-Wen Hu, Isabella Palazzo, Marisol Cano, Thanh Hoang, Imran Bhutto, Shusheng Wang, Debasish Sinha, Seth Blackshaw, Jiang Qian, James T. Handa

## Abstract

Environmental exposure such as cigarette smoke induces epigenetic changes that can induce degenerative heterogeneity and accelerate aging. In early age-related macular degeneration (AMD), the leading worldwide cause of blindness among the elderly, retinal pigment epithelial (RPE) cell heterogeneity is a key change. Since smoking is the strongest environmental risk factor for AMD, we hypothesized that cigarette smoke induces degenerative RPE heterogeneity through epigenetic changes that are distinct from aging, and that with aging, the RPE becomes vulnerable to cigarette smoke insult. We administered cigarette smoke condensate (CSC) intravitreally to young and aged mice and performed snRNA-seq and snATAC-seq on the RPE/choroid. This analysis identified separate cell clusters corresponding to healthy and abnormal, dedifferentiated RPE in both aged vehicle-treated and young CSC-treated mice. The dedifferentiated RPE were characterized by a global decrease in chromatin accessibility and decreased expression of genes in functional categories that were linked to hallmarks of aging. Notably, young, dedifferentiated RPE also exhibited a compensatory upregulation of hallmarks of aging-related genes, specifically those related to mitochondrial function and proteostasis. In contrast, aged dedifferentiated RPE did not express these compensatory changes, and did not survive CSC treatment, as experimentally verified with TUNEL labeling. These changes are relevant to early AMD because we identified through scRNA-seq, similar dedifferentiated and healthy macular RPE clusters in a donor who smoked and another with early AMD, but not from a nonsmoker. Degenerative cellular heterogeneity can include an abnormal cluster that jeopardizes cell survival and may represent an additional hallmark of ocular aging.

## Introduction

Aging was originally proposed to result directly and primarily from the accumulation of double-stranded DNA breaks^1,2^. Environmental stresses that promote DNA damage such as smoking can promote aging because it induces DNA damage^3^. However, mice deficient in the DNA mismatch repair gene *Pms2* accumulate mutations, and have normal health and lifespans, indicating that DNA damage alone is insufficient to drive aging.^4^ While DNA mutations clearly contribute to aging in multiple cell types,^5^ epigenetic changes have emerged as an additional mechanism for cellular aging.^6,7^ As a result, epigenetic clocks are now acknowledged as a marker of biological age in mammals^8,9^. In fact, epigenetic alterations are one of the twelve hallmark of aging-related changes that also includes autophagy dysfunction, cellular senescence, dysbiosis, genomic instability, inflammation, intercellular communication alterations, metabolism/nutrient-sensing dysregulation, mitochondrial dysfunction, proteostasis impairment, stem cell exhaustion, and telomere attrition^10–14^.

Epigenetic changes play an important role in cellular adaptation. In response to external stress, epigenetic changes induce long-term changes in subsequent responses to stimuli^15,16^. This adaptation is characteristic of ordered heterogeneity, a feature of any organism, which diversifies the cytoprotective response and is in part, regulated by the epigenome, which varies among individual cells^17^. In contrast, degenerative heterogeneity is disordered and compromises a tissue’s stress response. Not surprisingly, aging through epigenetic alterations is a mechanism of degenerative heterogeneity. When degenerative heterogeneity contains leads to the formation of variant cell states that can act in a non-cell autonomous manner to impair the function of healthy cells, aging can transition to disease states that include vascular pathologies, Alzheimer’s disease, and cancer^18–26^, even if only a few pathologic cells are present^27^. Age-dependent formation of senescent cells represent one such example^28,29^.

The retinal pigmented epithelium (RPE) is a critical, multifunctional cell that maintains the health of adjacent light sensing photoreceptor cells in the retina. Prior studies have found that the RPE displays ordered heterogeneity in the healthy, young eyes of multiple mammalian species^30–32^. Human eyes also display RPE heterogeneity, which is likely degenerative, given the advanced age of the donor eyes studied^33–40^. The RPE is a key cell type involved in the pathobiology of age-related macular degeneration (AMD), the most common global cause of irreversible vision loss among people over 60 years old^41^. In early-stage AMD, the RPE exhibits cellular heterogeneity^36,40,42–46^. RPE cells first undergo degeneration in the parafoveal macula, which is molecularly and morphologically distinct from foveal and peripheral RPE^47–52^. This heterogeneity indicates that a limited number of RPE cells are involved in the conversion from aging to early AMD^35,47–53^, and that its heterogeneity might be caused by a variable response to aging and/or environmental stresses. Cigarette smoking, the strongest environmental risk factor for AMD onset and progression^54–60^, in addition to inducing DNA damage, induces changes in DNA methylation and histone modifications, leading to changes in cellular epigenetic states^61–63^. Through epigenetic alterations, smoke can reprogram the transcriptome to simultaneously disrupt multiple cytoprotective pathways—a fundamental characteristic of a complex disease like AMD^61,64–71^. However, it is unclear how RPE degenerative heterogeneity develops and progresses with aging or smoking, or how it contributes to early AMD pathobiology.

If the intensity of external stresses surpasses a critical threshold, the induction of ordered heterogeneity will fail. With aging, this threshold is likely reduced and an impaired response may become permanent, triggering disease onset.^72^ At present, the extent that epigenetic changes from aging and smoking induce similar or different degenerative heterogeneity is unclear. Furthermore, it is unknown how vulnerable the aging RPE is to an external stimulus such as cigarette smoke. We hypothesized that cigarette smoke induces degenerative heterogeneity through epigenetic changes that is distinct from aging, and that with aging, the RPE are more vulnerable to cigarette smoke insult than young RPE. To address this hypothesis, we determined the extent that cigarette smoke induces epigenetic and transcriptomic changes in both young and aged mice and correlated these changes with human samples from a smoker and an early-stage AMD patient.

## Results

### Intravitreal cigarette smoke condensate (CSC) and aging induce two transcriptomically distinct RPE clusters

The earliest morphologic changes in aging and AMD occur in the parafoveal macula. Since the RPE and cone/rod density of mice is similar to the human parafovea, we chose mice as an *in vivo* model to study the impact of smoking and aging^73^. Three-month-old C57BL6J mice were given intravitreal (IVT) CSC or vehicle, and snRNA-seq and snATAC-seq were performed. We obtained 104,525 cells from young and aged mice after either IVT vehicle or CSC treatment. Nineteen clusters were identified, which includes all major cell types such as rod, cone, smooth muscle, and microglia (**Fig S1A, B**). A single RPE cluster was observed 3 days after either IVT vehicle or CSC based on the expression level of 15 RPE marker genes. Six and 10 days after IVT CSC, a second RPE cluster was identified, while mice given IVT vehicle maintained a single RPE cluster (**Fig 1A)**. Likewise, when 12-month-old mice were given either IVT vehicle or CSC, those given vehicle showed two distinct RPE clusters. This second cluster was designated as “dedifferentiated” because the expression of the core RPE gene set was decreased relative to the second cluster (**Fig 1B**). For example, *Slc6a20a* and *Myrip* were robustly expressed by healthy RPE, but were decreased in the dedifferentiated RPE cluster both following IVT CSC in 3-month-old mice or in IVT vehicle-treated 12-month-old mice (**Fig 1C).** The dedifferentiated RPE cluster was not observed in 12-month-old mice treated with IVT CSC **(Fig 1A**). The relative fraction of dedifferentiated RPE cells relative to healthy RPE cells increased following IVT CSC in young mice, eventually matching the relative fraction seen in vehicle-treated 12-month-old mice (**Fig 1C**). In contrast, choroidal cells did not resolve into separate cellular clusters either in response to CSC, which suggests that the RPE are uniquely vulnerable for developing stress-induced cellular heterogeneity (**Fig S1D**).

**Figure 1.**
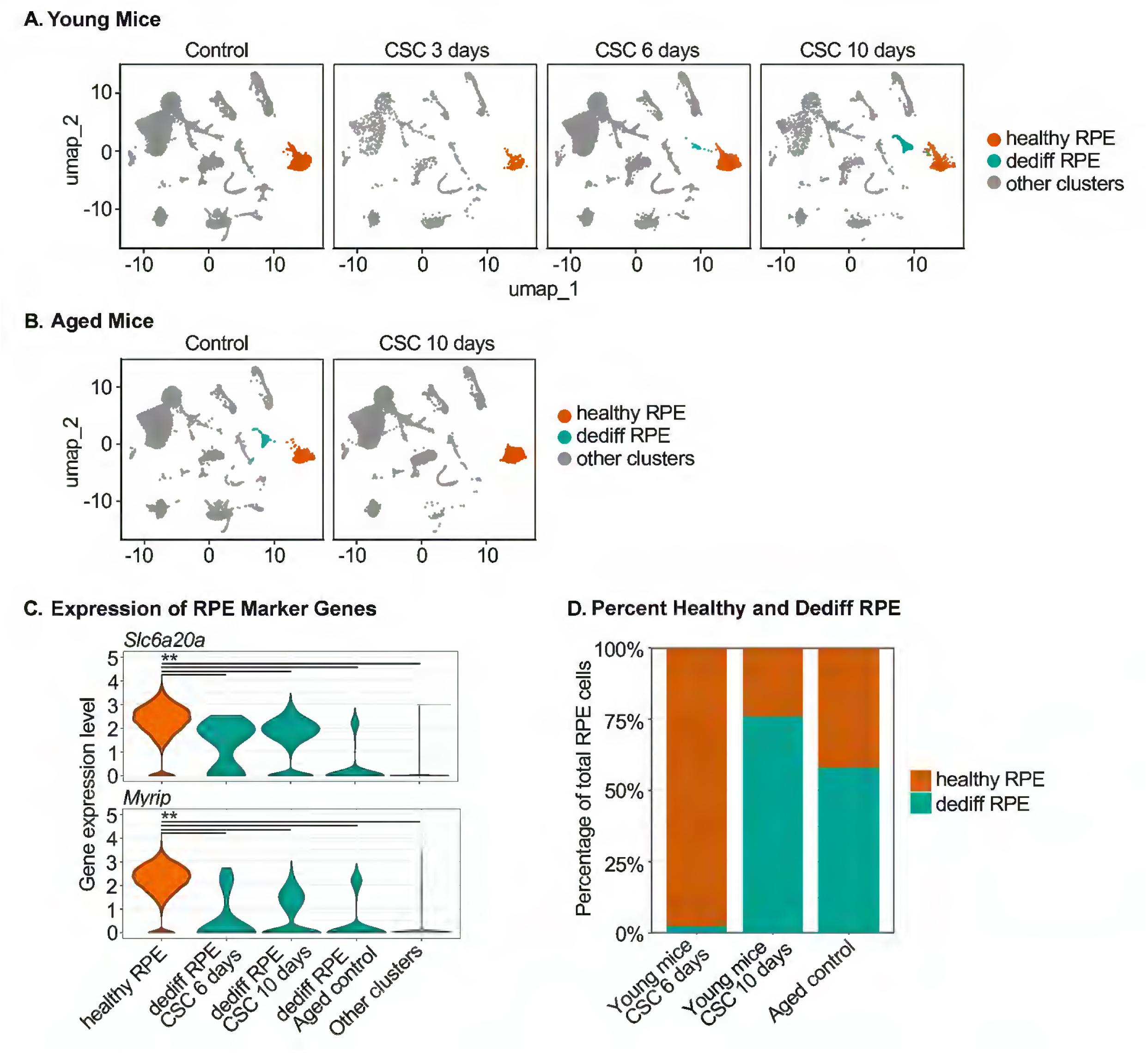
snRNA-seq of control and CSC-treated mice. **A)** snRNA-seq umaps of control, and after 3, 6, and 10 days following IVT CSC given to 3-month old mice, and **B)** Umap of 12-month old mice 10 days following IVT CSC. **C)** Expression of RPE marker genes *Slc6a20a* and *Myrip*. **D)** The percentage of dedifferentiated relative to healthy RPE cells increased with time following IVT CSC and with aging.

### Both CSC and aging induce two separate RPE clusters with different chromatin accessibility profiles

Previously, we observed decreased chromatin accessibility in the promoter regions of genes expressed in RPE cells of AMD maculas, which closely resembled changes observed in iPSC-RPE cells exposed to CSC^74^. To determine the extent that CSC or aging-induced decreases in chromatin accessibility influences RPE transcriptomic heterogeneity, snATAC-seq was performed on the RPE/choroid of young and aged mice given IVT CSC or vehicle. The RPE from 3-month-old eyes treated with vehicle had one chromatin accessibility cluster 3 days after CSC treatment and segregated into two clusters by 6 days following CSC (**Fig 2A**). The percentage of RPE cells in the reduced chromatin accessibility cluster increased by 10 days following CSC. In the RPE of aged mice, a normal and decreased chromatin accessibility cluster were also identified, although the decreased accessibility cluster disappeared following IVT CSC (**Fig 2B**). Unlike the decreased chromatin accessibility cluster observed in CSC-treated young mice, which contain almost only dedifferentiated RPE, the decreased chromatin accessibility cluster in aged mice also included other cell types. This observation suggests that aging may broadly reduce chromatin accessibility in multiple cell types, while CSC treatment predominantly affects RPE cells. Overall chromatin accessibility in the normal RPE cluster was reduced relative to choroidal cells and was further decreased in the dedifferentiated RPE cluster (**Fig 2C**). Similarly, the chromatin accessibility of aged RPE was lower overall than in choroidal cells, and further decreased in dedifferentiated RPE cells (**Fig 2D**). For example, the promoter regions of *Abhd2* and *Col9a3* showed decreased chromatin accessibility both in the young, dedifferentiated RPE observed following CSC treatment and in the aged, dedifferentiated RPE clusters relative to the vehicle-treated young RPE cluster (**Fig 2E**). Overall, chromatin accessibility decreased further by 10 days following IVT CSC, with 99% of all identified peaks reduced in dedifferentiated RPE. This reduction in chromatin accessibility resembled that observed in aged dedifferentiated RPE (**Fig 2F**), of which approximately 50% were in the gene promoters and correlated with decreased differentially expressed genes (DEGs) in dedifferentiated RPE induced by either CSC treatment or aging (**Fig 2G**).

**Figure 2.**
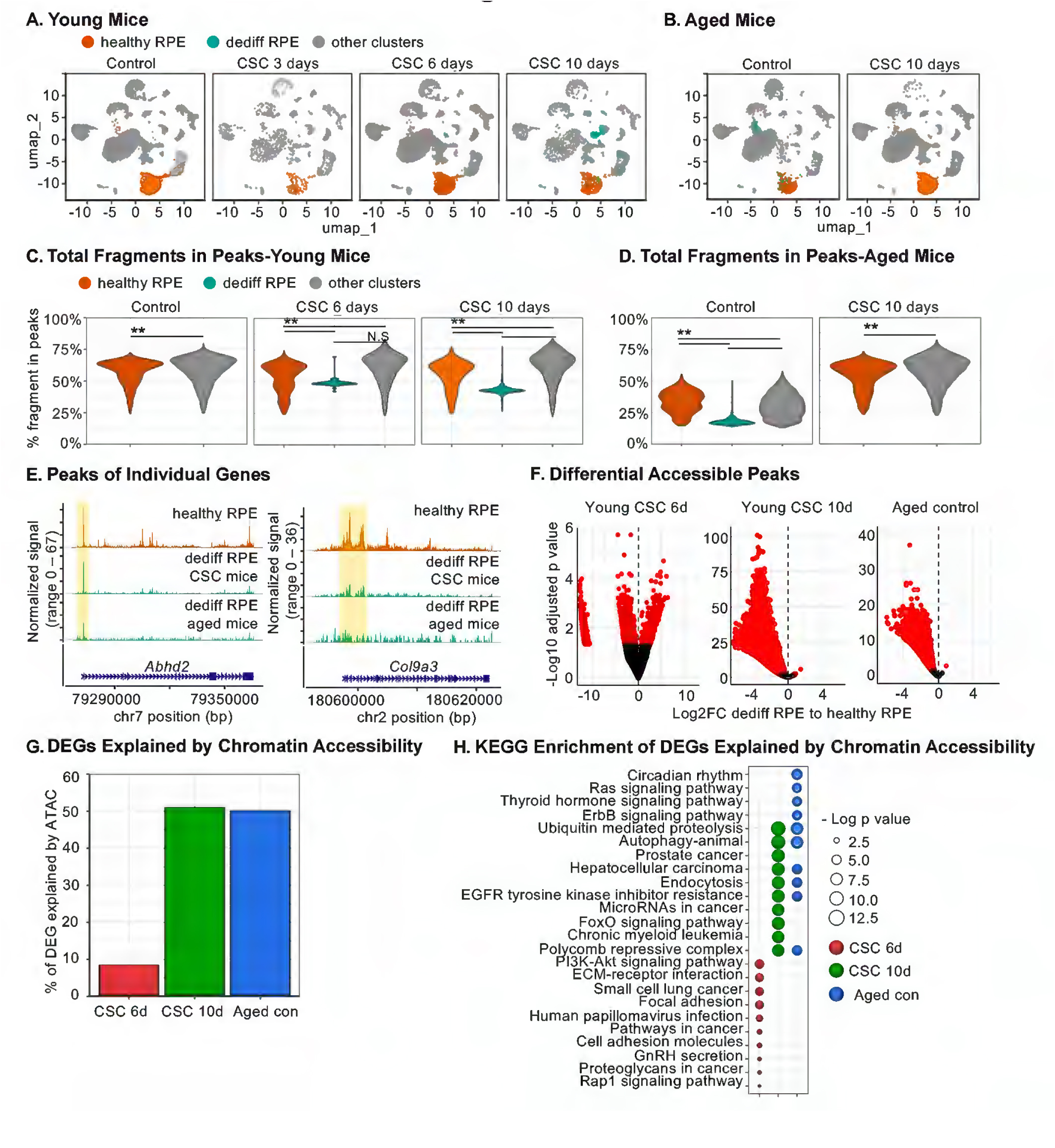
Chromatin accessibility changes of genes in the RPE following CSC and vehicle treatment. **A)** snATAC-seq umaps of control, and after 3, 6, and 10 days following IVT vehicle or CSC given to 3-month-old mice, and **B)** Umap of 12-month old mice 10 days following IVT vehicle or CSC. **C)** Total fragment in peaks of young healthy and dedifferentiated RPE, and choroidal cells following IVT vehicle or CSC at 6 and 10 days. **D)** Total fragment in peaks in aged healthy and dedifferentiated RPE, and choroidal cells following IVT vehicle or CSC after 10 days. **E)** Decline of chromatin accessibility peaks of *Abhd2* and *Col9a3* by position in dedifferentiated RPE. **F)** Volcano plot of differential accessibility peaks of genes in the RPE after either IVT CSC in young or vehicle in aged mice. **G)** Differentially expressed genes (DEGs) that have decreased chromatin accessibility in their promoter regions after IVT CSC or with aging. **H)** KEGG enrichment analysis DEGs with decreased chromatin accessibility in their promoter regions.

### The transcriptome of dedifferentiated RPE induced by CSC has similarities and differences with aging dedifferentiated RPE

The DEGs between the dedifferentiated and healthy RPE clusters reflect the changes to the dedifferentiated cell’s transcriptome. Using a Log2FC>0.1 and padj <0.05 cutoff, the number of DEGs between healthy RPE in vehicle and CSC-treated eyes was low. In contrast, the number of down-or upregulated DEGs between dedifferentiated and healthy RPE steadily increased from 6 to 10 days following CSC treatment (**Fig 3A**). Relative to healthy RPE, dedifferentiated RPE showed 52% downregulated and 48% upregulated DEGs at 6 days, and 61% downregulated and 39% upregulated DEGs at 10 days following CSC treatment. In aged animals, dedifferentiated RPE had 75% downregulated and 25% upregulated DEGs (**Fig 3B**). Distinct differences were seen in the transcriptional response to CSC and aging. When the expression of genes in young, dedifferentiated CSC-treated RPE at 10 days was compared to aged, dedifferentiated vehicle-treated RPE, the majority of DEGs were upregulated (**Fig 3A**), broadly reflecting observed changes in chromatin accessibility.

**Figure 3.**
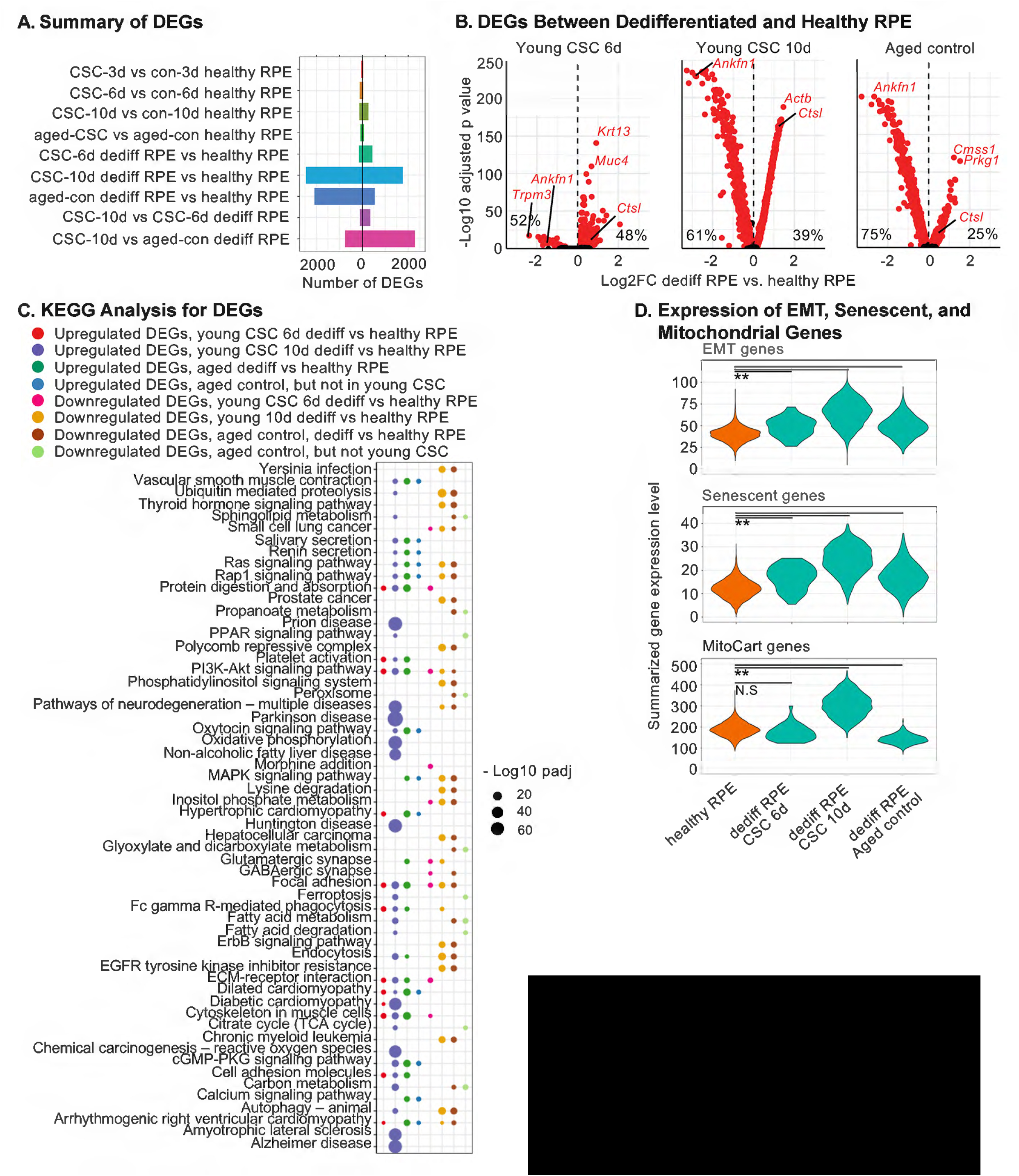
The Transcriptional phenotype of dedifferentiated RPE following CSC treatment and with aging. **A)** Up and downregulated DEGs in dedifferentiated relative to healthy RPE after IVT CSC or with aging. **B)** Volcano plots of up and downregulated DEGs in dedifferentiated RPE after IVT CSC or with aging. Note how *Ankfn1* is markedly decreased and *Ctsl* is increased 10 days following CSC-treated young and vehicle-treated aged dedifferentiated RPE. **C)** KEGG enrichment analysis of DEGs in dedifferentiated RPE following IVT CSC or with aging. **D)** Expression of EMT and mito-senescent genes from the Broad Institute’s EMT gene set and SenMayo gene set^78^, respectively, and mitochondrial genes from the Broad Institute’s Mito-carta V3 gene set in the RPE following IVT CSC or with aging.

KEGG enrichment analysis was used to compare dedifferentiated versus healthy RPE to analyze transcriptional changes linked to decreased chromatin accessibility following CSC treatment (**Fig 2H, 3C**). DEGs were analyzed at 10 days post-CSC because the most highly downregulated DEGs at 6 days post-CSC were not sustained at later timepoints. Most KEGG pathways enriched with downregulated DEGs in young, CSC-treated dedifferentiated RPE were also enriched in 8 of the 12 functional categories identified as hallmarks of aging, which include autophagy dysfunction, cellular senescence, epigenetic alterations, genomic instability, inflammation, metabolism/nutrient-sensing dysregulation, mitochondrial dysfunction, and proteostasis impairment (**Fig 2H**; **Table 1; Table S1, Fig S2)**.^10–14^ Likewise, KEGG pathways that were selectively enriched in downregulated DEGs in dedifferentiated RPE of aged vehicle-treated mice, were also enriched in these same seven hallmarks of aging categories, with the sole exception of epigenetic alterations (**Fig 2H**; **Table 1; Table S3, Fig S4**).^10–14^ Whether due to CSC stress or aging, these hallmarks of aging categories often involved genes in the MAPK and PI3K-AKT pathways.

**Table 1.** DEGs in dedifferentiated RPE in KEGG pathways that are associated with hallmark of aging-related changes and the MAPK/PI3K-AKT pathways.

| Table 1 |  |  |  |
| --- | --- | --- | --- |
| CSC Treated Mice |  |  |  |
| # Decreased DEGs | KEGG Pathway | Hallmark Aging Category | # MAPK/PI3K-AKT genes |
| 43 | Autophagy | Autophagy impairment | 27 |
| 24 | Chronic myeloid leukemia | Genomic instability | 17 |
| 25 | EGFR tyrosine kinase inhibitor resistance | Genomic instability, mitochondrial dysfunction, immune dysregulation, proteostasis impairment | 25 |
| 53 | Endocytosis | Immune dysregulation, proteostasis impairment | 5 |
| 33 | FoxO signaling pathway | Autophagy impairment, epigenetic alterations, genomic instability, immune dysregulation, metabolism impairment, proteostasis impairment | 25 |
| 41 | Hepatocellular carcinoma | Cellular senescence, epigenetic alterations, genomic instability, proteostasis impairment | 30 |
| 38 | MicroRNAs in cancer | Epigenetic alterations, mitochondrial dysfunction | 21 |
| 25 | Polycomb repressive complex | Epigenetic alterations | 0 |
| 30 | Prostate cancer | Epigenetic changes, genomic instability | 28 |
| 41 | Ubiquitin mediated proteolysis | Proteostasis impairment | 2 |
| # Increased DEGs | KEGG Pathway | Hallmark Aging Category | # MAPK/PI3K-AKT genes |
| 107 | Alzheimer disease | Cellular senescence, metabolism impairment, mitochondrial dysfunction, proteostasis impairment | 3 |
| 99 | Amyotrophic lateral sclerosis | Mitochondrial impairment, proteostasis impairment | 2 |
| 72 | Chemical carcinogenesis - reactive oxygen species | Autophagy impairment, mitochondrial dysfunction, proteostasis impairment | 1 |
| 69 | Diabetic cardiomyopathy | Autophagy impairment, cellular senescence, metabolism impairment, mitochondrial dysfunction, proteostasis impairment | 0 |
| 95 | Huntington disease | Autophagy impairment, mitochondrial dysfunction, proteostasis impairment | 0 |
| 58 | Non-alcoholic fatty liver disease | Cellular senescence, metabolism impairment, mitochondrial dysfunction | 2 |
| 62 | Oxidative phosphorylation | Mitochondrial dysfunction | 0 |
| 106 | Parkinson disease | Mitochondrial dysfunction, proteostasis impairment | 1 |
| 118 | Pathways of neurodegeneration - multiple diseases | Autophagy impairment, immune dysregulation, metabolism impairment, mitochondrial dysfunction, proteostasis impairment | 4 |
| 93 | Prion disease | Autophagy dysfunction, immune dysregulation, mitochondrial dysfunction, proteostasis impairment | 1 |
| Aged Mice |  |  |  |
| # Decreased DEGs | KEGG Pathway | Hallmark Aging Category | # MAPK/PI3K-AKT genes |
| 30 | Autophagy - animal | Autophagy dysfunction | 17 |
| 11 | Circadian rhythm | Proteostasis impairment | 0 |
| 16 | EGFR tyrosine kinase inhibitor resistance | Genomic instability, immune dysregulation, mitochondrial dysfunction, proteostasis impairment | 16 |
| 33 | Endocytosis | Immune dysregulation | 2 |
| 16 | ErbB signaling pathway | Autophagy impairment, genomic instability, metabolism impairment, proteostasis impairment | 12 |
| 26 | Hepatocellular carcinoma | Cellular senescence, epigenetic changes, genomic instability, proteostasis impairment | 18 |
| 16 | Polycomb repressive complex | Epigenetic alterations | 0 |
| 31 | Ras signaling pathway | Cellular senescence | 26 |
| 20 | Thyroid hormone signaling pathway | Epigenetic alterations, immune dysregulation, proteostasis impairment | 11 |
| 28 | Ubiquitin mediated proteolysis | Proteostasis impairment | 0 |
| # Increased DEGs | KEGG Pathway | Hallmark Aging Category | # MAPK/PI3K-AKT genes |
| 15 | Calcium signaling pathway | Cellular senescence, metabolism impairment, mitochondrial dysfunction | 1 |
| 18 | cGMP-PKG signaling pathway | Cellular senescence, mitochondrial dysfunction | 0 |
| 14 | Dilated cardiomyopathy | Cellular senescence/ telomere shortening | 0 |
| 14 | MAPK signaling pathway | Cellular senescence | 14 |
| 16 | Rap1 signaling pathway | Cellular senescence | 0 |

The key difference between young, dedifferentiated CSC-treated RPE and aged, vehicle-treated dedifferentiated RPE was upregulation of DEGs enriched in KEGG pathways involved in five hallmarks of aging categories, especially mitochondrial function and proteostasis, and to a lesser extent autophagy, inflammation, and metabolism (**Fig 3C**, **Table 1, Table S2, Fig S3)**.^10–14^ While some upregulated DEGs were observed in dedifferentiated RPE from vehicle-treated aged mice, the overall increase in expression was modest (**Table 1; Table S4; Fig S5**).

### Evidence of EMT, mito-senescent, and mitoCarta (3.0) phenotype in dedifferentiated RPE

We previously showed that RPE cells with mitochondrial dysfunction undergo epithelial-mesenchymal transition (EMT) through retrograde mitochondrial to nuclear signaling^75^. Yu *et al.* have likewise shown that the RPE can become senescent^76^. Cellular senescence, a hallmark of aging^10–14^, can be induced by different triggers including mitochondrial dysfunction-associated senescence (MiDAS)^77^. Given the enrichment of DEGs involved in mitochondrial function and the possibility of MiDAS, we next evaluated for differentially expressed mitochondrial genes. Using the Broad Institute’s MitoCarta 3.0 inventory of 1136 human and 1140 mouse genes encoding proteins with mitochondrial function, we found that compared to healthy RPE, dedifferentiated RPE from both young CSC-treated and aged vehicle-treated mice showed altered expression of mitochondrial genes (**Fig 3D**). Notably, mitochondrial genes were upregulated in young, dedifferentiated CSC-treated RPE, while their expression was decreased in aged vehicle-treated RPE. We next analyzed dedifferentiated RPE for evidence of EMT using the Broad Institute’s EMT gene set and, given the enrichment of differentially expressed mitochondrial genes, for mito-senescence using the SenMayo gene set^78^. In both young mice treated with IVT CSC and aged mice, dedifferentiated RPE expressed a mixture of both EMT and senescent transcriptional profiles, and thus, is its own unique phenotype (**Fig 3D**).

### The distribution and morphology of dedifferentiated and healthy RPE

We next sought to identify the distribution of dedifferentiated RPE in the fundus. As shown in **Fig 3B**, because *Ankfn1* was abundant in healthy RPE, but markedly decreased in dedifferentiated RPE, we selected it a marker for healthy RPE. With markedly increased expression in dedifferentiated RPE and minimal expression in healthy RPE, *Ctsl* was chosen to label dedifferentiated RPE. Using RNAscope on 3-month-old mice given either IVT DMSO or CSC, we found that *Ankfn1*-expressing healthy RPE were distributed throughout the fundus of DMSO-treated mice (**Fig 4A-F**) while *Cstl1*-expressing dedifferentiated RPE of CSC-treated mice were enriched in the posterior relative to anterior fundus (**Fig 4G-L**). While *Ankfn1*-labeled RPE had cobblestone morphology, *Cstl1*-labeled dedifferentiated RPE were enlarged and elongated relative to healthy cells (**Fig 4M,N**).

**Figure 4.**
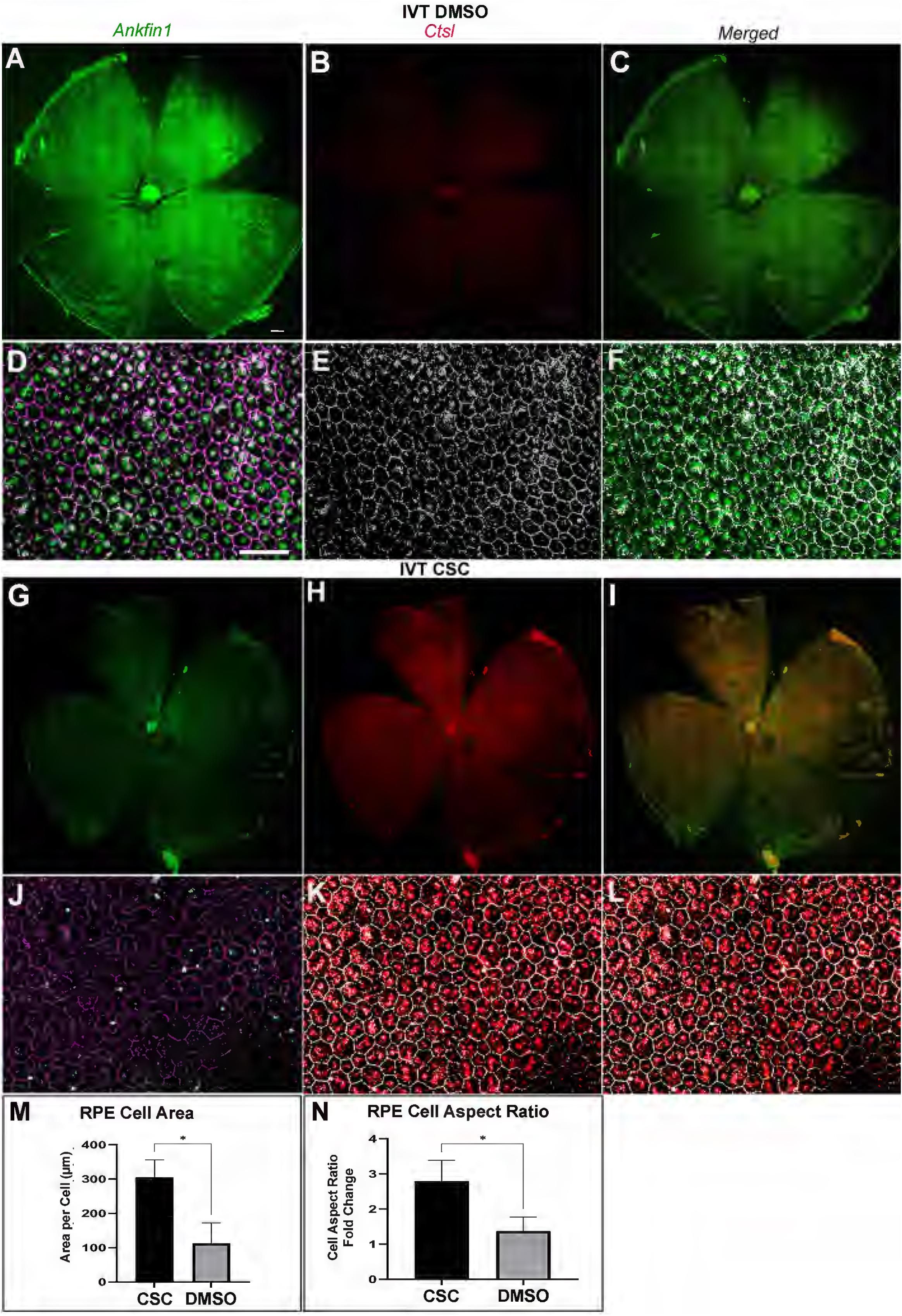
Morphology and distribution of healthy and dedifferentiated RPE after IVT CSC. RPE flatmounts from a 3-month old mouse showing the distribution of healthy RPE labeled with *Ankfn1* (green) and dedifferentiated RPE labeled with *Ctsl* (red) ten days after either IVT DMSO or CSC. **A)** Low power image of whole RPE flatmount of a mouse given IVT DMSO shows predominantly *Ankfn1* (green) labeled healthy RPE. **B)** Same flatmount with few RPE labeled for *Ctsl* (red) in dedifferentiated RPE. **C)** Merged image showing predominantly *Ankfn1* labeled RPE. High power view of **D)** *Ankfn1*, **E)** *Ctsl*, and **F)** merged *Ankfn1* and *Ctsl* labeled images with ZO-1 (magenta in D, falsely colored white in E, F) immunolabeling to show RPE cell shape. Low power image of whole RPE flatmount of a mouse given IVT CSC. **G)** *Ankfn1* (green) labeled healthy RPE are fewer in number compared to DMSO treated eyes. **H)** *Ctsl* (red) labeled dedifferentiated RPE in high number. **I)** Merged image of *Ankfn1* and *Ctsl* labeled RPE. High power image of **J)** *Ankfn1*, **K)** *Ctsl*, and **L)** merged *Ankfn1* and *Ctsl* labeled images with ZO-1 (falsely colored white in K, L) immunolabeling to show RPE cell shape. Bar=1500 µm for low power images and 50 µm for high power images. **M, N)** Graph of cell area and cell aspect ratio, respectively, as calculated in^75^, *p<0.05.

### Dedifferentiated RPE cells from aged mice die by apoptosis

In 12-month aged mice, the dedifferentiated, transcriptional RPE cluster was lost by ten days following IVT CSC. With KEGG analysis, we found that aging, dedifferentiated RPE were enriched with DEGs related to multiple cell death-related pathways, including apoptosis (**Fig 5A**). Since RPE cells can die by apoptosis in both natural aging and AMD^79,80^, we explored if apoptosis was induced following CSC stress. Three– and 12-month-old mice were given IVT CSC. After 72 hours, dedifferentiated RPE of 3-month-old mice given IVT CSC were labeled with *Ctsl1* mRNA with minimal TUNEL labeling (**Fig 5B**). In contrast, 12-month-old mice given IVT CSC have dedifferentiated RPE that co-label with both *Ctsl1* mRNA and TUNEL signal, indicating that dedifferentiated RPE cells die by apoptosis in aged animals while 12-month-old mice given IVT DMSO have dedifferentiated RPE that label with *Ctsl1* mRNA but show minimal TUNEL signal (**Fig 5C**).

**Figure 5.**
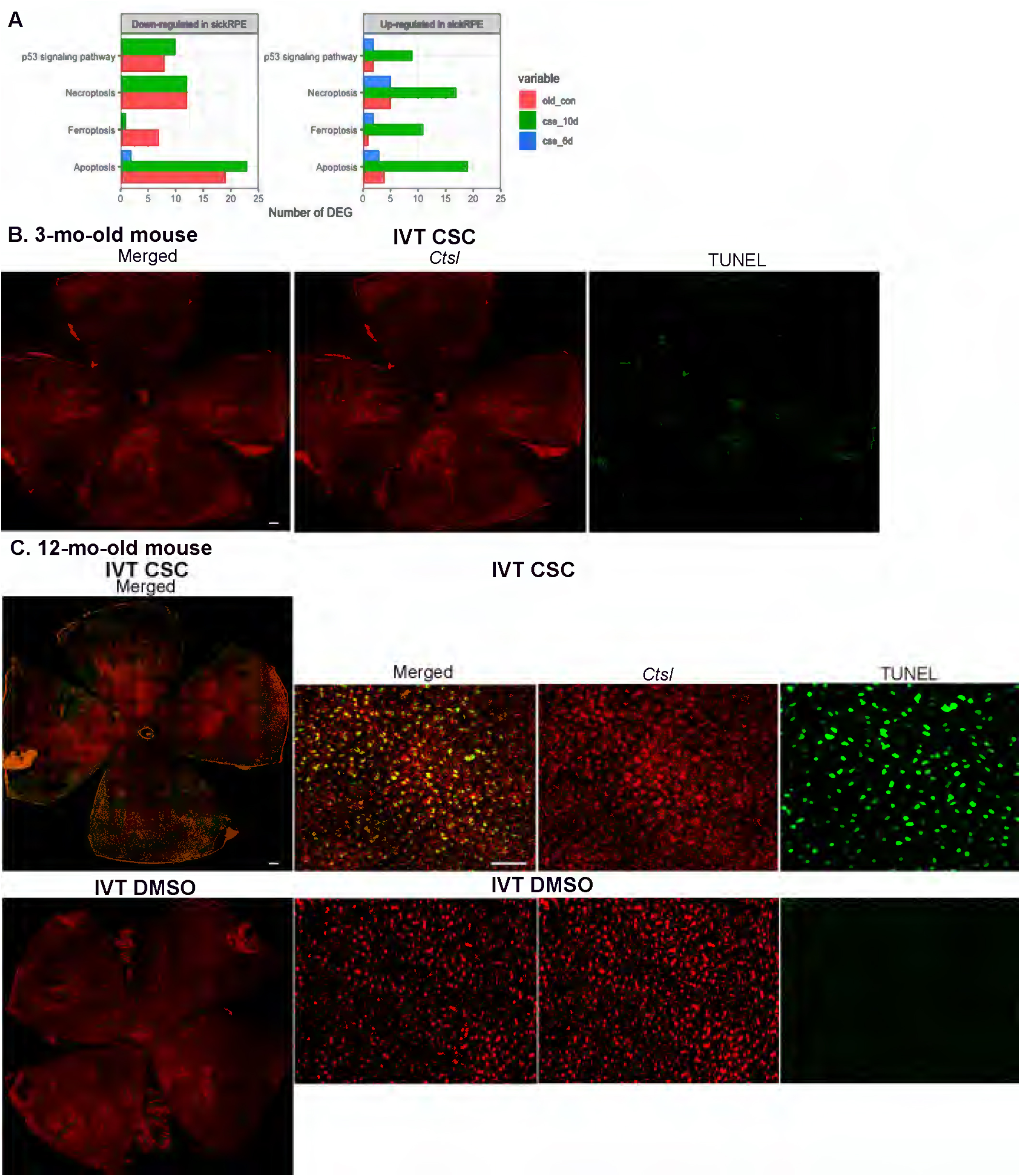
Dedifferentiated RPE die by apoptosis. **A)** KEGG enrichment death associated pathways of down– and up-regulated DEGs in dedifferentiated RPE following IVT CSC and with aging. **B)** 3-month old mouse given IVT CSC. Low power image of whole RPE flatmount. Left shows merged RNAscope labeling of *Ctsl* (red) and TUNEL (green), which shows minimal cell death. Bar=1500 µm **C)** 12-month old mouse given IVT CSC (top) or DMSO (bottom). Top left shows low power of whole RPE flatmount of merged, *Ctsl* (red), and TUNEL (green) labeling. Bar=1500 µm. Top right are high power views of merged, *Ctsl* and TUNEL labeling. Bottom left shows low power RPE flatmount with abundant RPE labeled for *Ctsl* with minimal TUNEL. Bottom right are high power views of merged, *Ctsl* and TUNEL labeling. Bar=50 µm.

### RPE from human macula segregates into two clusters with smoking and early AMD

These findings raise the possibility that similar changes may occur in human RPE cells obtained from a donor without AMD who smoked or one with early-stage AMD. Using the 154 genes identified by Srinivas *et al.* to identify human RPE cells^81^, healthy and dedifferentiated RPE cell clusters from the macula were identified in a smoker without AMD (smoker-control) and an early AMD donor (Minnesota Grade 2), while the RPE cells from the nonsmoker donor without AMD (nonsmoker control) formed a single cluster (**Fig 6A**). *RPE65* and *OTX2* were markedly decreased in dedifferentiated relative to healthy RPE cells from the nonsmoker-control (**Fig 6B, C**). The percentage of macular RPE in the dedifferentiated cluster was increased in the early AMD donor relative to the control-smoker donor (**Fig 6C**).

**Figure 6.**
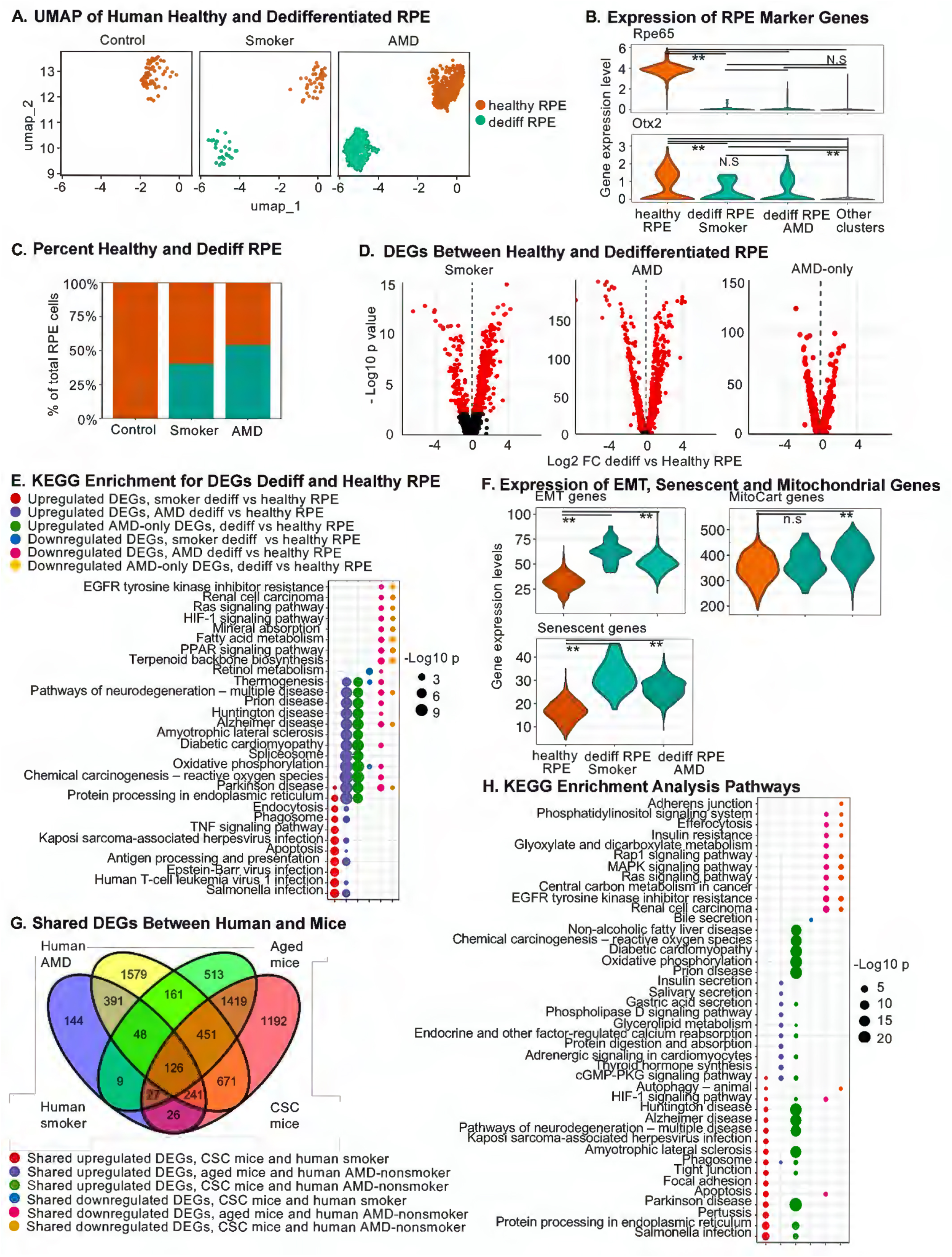
The Transcriptional phenotype of macular RPE from a non-smoker control, smoker control, and early AMD donor. **A)** snRNA-seq umaps of nonsmoker-control, smoker-control, and early AMD macular RPE showing two RPE populations in the smoker-control and early AMD donor eyes. **B)** Decreased expression of RPE marker genes *RPE65* and *OTX2*. **p<0.01**. C)** The % dedifferentiated RPE is increased in the early AMD macula compared to the smoker-control eye. **D)** Volcano plots of up and downregulated DEGs in the smoker-control and early AMD. In addition, the distribution of DEGs in the AMD macular RPE (AMD-only) is shown after removing DEGs that were in common with the smoker-control. **E)** KEGG enrichment analysis of DEGs in the dedifferentiated versus healthy RPE between smoker-control and early AMD including AMD-only. **F)** Expression of EMT, senescence, and mitoCarta genes in healthy and dedifferentiated macular RPE. **p<0.01. **G)** Venn diagram of shared DEGs between human macular RPE and mouse RPE. **H)** KEGG enrichment analysis of shared DEGs in dedifferentiated RPE between young mouse RPE treated with CSC or aged mouse RPE and early AMD or AMD-only.

We reasoned that up-or down-regulated DEGs between dedifferentiated and healthy RPE in the AMD donor after excluding DEGs observed in the smoker-control, suggested a link with AMD (AMD-only). In contrast, we expect that DEGs observed between the dedifferentiated and healthy RPE from the smoker-control, but not the AMD donor, would be associated with smoking and not AMD. Using this criteria, we identified up– and down-regulated DEGs, as illustrated in the Volcano plots (**Fig 6D**). We next identified DEGs enriched in KEGG pathways that were associated with AMD-only (**Fig 6E**). The top KEGG pathways that were enriched exclusively with upregulated DEGs included amyotrophic lateral sclerosis and the spliceosome. In addition, nine other KEGG pathways with upregulated DEGs included modestly downregulated DEGs. These eleven KEGG pathways were enriched with upregulated DEGs involved in hallmark of aging categories including autophagy, metabolism, mitochondrial function, and proteostasis (**Fig S7**).

KEGG pathways enriched with downregulated DEGs in dedifferentiated RPE of the AMD-only donor were also identified, which were linked with hallmark of aging-related changes including metabolism dysregulation, mitochondrial dysfunction, and proteostasis impairment. Downregulated DEGs in the MAPK and PI3K-AKT signaling pathways were either linked with these hallmarks of aging-associated changes or involved with cell survival, proliferation, and differentiation (**Fig S8**). Dedifferentiated RPE from the AMD-only donor were enriched for increased expression of genes related to EMT, senescence, and MitoCarta (**Fig 6F**). Thus, the transcriptional profile of dedifferentiated RPE in early AMD is a unique phenotype with a mixed induction and decline of genes involved with hallmarks of aging-associated categories including mitochondrial function, metabolism, and proteolysis.

### Similarity of dedifferentiated RPE transcriptional phenotype of mouse and human

1698 DEGs (24%) in dedifferentiated RPE from human maculas overlapped with DEGs in dedifferentiated RPE cells from both CSC-treated young mice or vehicle-treated aged mice (**Fig 6G**). Specifically, 1489 DEGs (21%) between dedifferentiated RPE cells from the AMD donor and young mice given IVT CSC, and 376 DEGs (5%) were shared between dedifferentiated RPE cells in AMD patients and aging mice. The phenotype for shared DEGs between the RPE of human and mice was characterized with KEGG enrichment analysis. The most significant upregulated DEGs in dedifferentiated RPE in both young CSC-treated mice and AMD patients were enriched in five KEGG pathways that principally involved mitochondrial function (**Fig 6H, Fig S9**). Upregulated DEGs observed in dedifferentiated RPE in aged vehicle-treated mice and AMD-only were enriched in multiple KEGG pathways that were related to small molecule transporters, although the enrichment was less significant as compared to the young CSC-treated and AMD-only RPE (**Fig 6H**).

Downregulated DEGs were shared between young, CSC-treated and AMD-only RPE in 9 KEGG pathways and were enriched in hallmarks of aging-related categories including autophagy impairment, metabolism/nutrient-sensing dysregulation, mitochondrial dysfunction, and proteostasis impairment (**Fig 6H, Fig S10**). Again, these pathways often involved the PI3K-AKT and MAPK signaling pathways that are linked with cell survival, proliferation, and differentiation (**Fig S10**). Downregulated DEGs in dedifferentiated RPE in common between vehicle-treated aged mice and AMD-only involved ten KEGG pathways, of which 8 pathways were also seen in young, dedifferentiated CSC-treated RPE (**Fig 6H**).

These pathways were enriched for the hallmark of aging-related changes including metabolism impairment, mitochondrial dysfunction, and proteostasis impairment, as well as cell survival pathways through the PI3K-AKT and MAPK signaling pathways (**Fig S11**). Thus, both young CSC-treated and aged vehicle-treated RPE shared with the early AMD donor a pattern of decreased expression of hallmark of aging-related genes that were associated with autophagy dysfunction, metabolism impairment, mitochondrial dysfunction, and proteostasis impairment that contain genes in the MAPK and PI3K-AKT pathways, along with induction of genes related to mitochondrial function.

## Discussion

The RPE is designed to keep photoreceptors healthy so they can process light for vision. The RPE apical microvilli extend to surround cone outer segments, but are shorter around rod outer segments^82^. This morphologic difference is an example of ordered heterogeneity, an adaptation that optimizes the RPE’s ability to maintain photoreceptor health. In contrast, we find that either aging or acute CSC stress in young mice induced RPE degenerative heterogeneity with an abnormal, dedifferentiated transcriptomic cluster that was linked with decreased chromatin accessibility. The dedifferentiated transcriptome was a unique phenotype comprised of a combination of both EMT and mito-senescent DEGs.

Importantly, the dedifferentiated cluster had decreased expression of genes in ‘hallmark of aging’ categories, which often involved the MAPK and PI3-AKT pathways. However, a crucial difference in the dedifferentiated RPE transcriptome was that young, but not aged, mice exposed to CSC were able to generate a compensatory response in some genes involved in functional categories related to established hallmarks of aging, especially mitochondrial function and proteostasis, which correlated with cell survival following CSC stress. In contrast, dedifferentiated RPE from aged mice did not exhibit these adaptive changes and did not survive an acute CSC challenge, as experimentally demonstrated with TUNEL labeled dedifferentiated RPE in aged, but not young CSC-treated mice. Finally, the heterogeneity induced by CSC treatment in young mice was unique to the RPE, while choroidal cell heterogeneity was also observed in aged mice. These findings identify mechanisms underlying the selective vulnerability of RPE cells to cellular stressors. Their relevance to human disease is underscored by the observation of a similar degenerative RPE transcriptomic heterogeneity profile in the macula of a smoker without AMD and a smoker with early AMD that included altered expression of hallmark of aging-related genes.

Aging results from the progressive, adverse influence of environmental stresses that jeopardize cellular homeostasis and cytoprotective responses. Twelve ‘hallmarks of aging’ have been proposed that include autophagy dysfunction, cellular senescence, dysbiosis, epigenetic alterations, genomic instability, inflammation, intercellular communication alterations, metabolism/nutrient-sensing dysregulation, mitochondrial dysfunction, proteostasis impairment, stem cell exhaustion, and telomere attrition^10–14^. These aging changes are interconnected with one another. For example, proteostasis impairment can induce senescence, which can be prevented by inhibiting mitochondrial respiration^83^. On the other hand, mitochondrial injury impairs proteostasis^84,85^. Interestingly, we found that epigenetic alterations, as measured by decreased chromatin accessibility in the promoter region of genes, was linked with their decreased expression in the dedifferentiated, but not healthy RPE cluster. This abnormal profile included decreased DEGs involved in seven hallmark of aging-related changes. Importantly, cellular heterogeneity can itself be driven by epigenetic changes^10,11,86^ where a small cell subpopulation can act in a non-cell autonomous manner to drive aging and pathological changes^18–27^. The cellular heterogeneity that is observed here may precede and underpin the formation of dedifferentiated cells, and itself is a potential hallmark of aging.

Among the hallmark of aging-related changes, genes in the MAPK and PI3K-AKT signaling networks were downregulated in dedifferentiated RPE of aging mice and CSC-treated young mice. The decreased expression of MAPK and PI3K-AKT pathway genes, which both influence cell proliferation, survival, and differentiation,^87–89^ was linked with decreased chromatin accessibility, which suggests that these cell survival pathways become impaired with aging and smoking through an epigenetic mechanism. These signaling networks were in turn, connected with other signaling pathways including calcium, mTOR, WNT, Notch, and NF-kB signaling, which in turn modulate hallmark of aging-related changes in functional categories that included autophagy dysfunction, genomic instability, inflammation, metabolism/deregulated nutrient-sensing impairment, and proteostasis impairment.

The MAPK and PI3K-AKT pathways are also involved with the development of EMT and senescence^90,91^. With EMT, epithelial cells develop a mesenchymal phenotype with enhanced ability to proliferate, and produce extracellular matrix with altered cell-cell adhesion and cell-matrix adhesion to promote migration, in an attempt to survive an insult^90^. In contrast, senescence is a program that permanently arrests proliferation with development of a senescence-associated secretory phenotype (SASP) that contributes to impaired tissue regeneration^91^. Since EMT and senescence are distinct cellular phenotypes, it is unusual that dedifferentiated RPE expressed elements of both programs. With additional stress, it is possible that the dedifferentiated RPE cluster will evolve into a distinct phenotype.

Experiments that interrogate the specific role of these signaling pathways are needed to define their contribution to EMT, senescence, and aging.

The decreased chromatin accessibility that was associated with decreased gene expression in the dedifferentiated RPE cluster of young mice suggests that besides aging, a specific environmental influence such as smoking can drive degenerative heterogeneous epigenetic changes that accelerates aging. The development of heterogeneity following CSC treatment in young mice was unique to the RPE and was more generalized with aging to involve choroidal cells. In addition, young dedifferentiated RPE cells—in contrast to the aged, dedifferentiated cells—upregulated genes associated with multiple hallmark of aging-related categories, particularly mitochondrial function and proteostasis, that are known to promote cellular homeostasis and viability^92^. The minimal levels of RPE cell death that are seen following CSC treatment in young mice suggests that the increased expression of hallmarks of aging-related genes in these cells was homeostatic and cytoprotective. On the other hand, the decreased expression of these genes in aged mice, possibly due to globally decreased chromatin accessibility, resulted in dedifferentiated RPE in aged mice that were unable to survive an acute CSC insult, as suggested experimentally by the TUNEL labeled *Ctsl* labeled cells, despite a mild induction of some hallmark of aging-related genes. Thus, the magnitude of decline in these hallmark of aging-related genes may determine the cell’s ability to maintain its homeostatic function and viability.

Multiple studies of elderly eyes have identified morphological differences between macular and peripheral RPE cells^33–40,46–48,50–52,93–100^. These differences correlate with transcriptomic changes between macular and peripheal RPE and indicate that with aging, the RPE develops degenerative heterogeneity that varies based on its spatial location. Our scRNA-seq data from human maculas differ from prior studies as we identified heterogeneity within the macula, linked this heterogeneity with smoking, and observed it in early AMD. The overall photoreceptor composition of the mouse retina resembles that of the rod-rich human parafovea^39,73^. Our findings in mice are therefore likely most relevant to the human parafovea, which is the location of the earliest AMD changes, including the onset of cellular heterogeneity in RPE cells^36,40,42–52^.

We recognize that our model may not simulate aging or the changes induced by long-term smoking. We acknowledge that the small sample size of high-quality human globes limits the interpretation of our findings to early AMD. Furthermore, our early AMD results may not be relevant to prior studies of advanced AMD because the pathogenic pathways in early disease are likely different from those seen in late-stage disease^92,98,101–103^. Using bulk RNA– and ATAC-seq analysis, we previously observed that reduced chromatin accessibility in promoter regions of genes occurred prior to transcriptional changes of the RPE in early AMD, and that the chromatin accessibility profile of iPSC-RPE cells exposed to CSC was similar to that of the RPE from AMD donors^74^. However, we do not know how much our transcriptional changes in the human eyes were induced epigenetic changes resulting from cellular heterogeneity, since we did not perform scATAC-seq analysis of these samples. While we did not genotype the human globes, we found in prior work that epigenetic changes were independent of the known genetic risk variants^74^.

Other studies have also reported cellular heterogeneity among RPE cells in both mice and humans. For instance, Lee *et al.* found 4 subpopulations of healthy RPE from similarly aged C57BL6 mice^104^. This group used different criteria to define an RPE cell. Similarly, Mullin *et al.* identified 4 RPE subpopulations in a healthy human donor eye based on expression levels of visual cycle genes^97^. However, it will be important in future experiments to standardize the definition of a normal RPE cell. While our cell death experiments imply the existence of two distinct cell populations in aged mouse RPE, future experiments must examine functional differences among the RPE subtypes to ultimately define the number of RPE subpopulations, and their role in aging and AMD.

Our studies find that the RPE of aged mice and young mice subject to acute CSC stress display degenerative heterogeneity that appears to be driven significantly by epigenetic changes that affect hallmark of aging-related changes, which was similar to the RPE heterogeneity from the macula of human smokers with or without early AMD. Notably, the dedifferentiated RPE of young mice induced the expression of some genes characteristic of hallmarks of aging, whereas these genes were downregulated in aged mice, which was correlated with induction of apoptosis following acute CSC stress. Future studies will explore how smoking and aging both induce epigenetic changes that lead to the formation of dedifferentiated RPE cells, and how this might in turn drive AMD pathogenesis.

## Materials and Methods

### Mouse care and IVT CSC protocol

Experiments were in accordance with NIH guidelines and approved by the Johns Hopkins Animal Care and Use Committee. C56BL/6J mice that were free of the *Rd8* mutation (Jackson Labs, Bar Harbor, ME) were given water ad libitum and kept in a 12-hr light–12-hr dark cycle. For experiments, an equal number of male and female mice aged either 3-or 12-months old were given an IVT injection of 250 µg/ml CSC in 1 μl (Murty Pharmaceuticals, Inc. Lexington, KY) or an equal volume (1 μl) of vehicle (DMSO) in both eyes using a microinjection pump (Harvard Apparatus, Holliston, MA, USA) on day 1 and day 3.

### Human tissue

The tenets of the Declaration for Helsinki for research involving human tissue were followed. Human eyes (n=3) were obtained from the Advancing Sight Network (Birmingham, AL). Preliminary experiments were conducted to determine the viability of RPE that was based on death to enucleation (D-E) time. Single cell RNA-seq on normal eyes from two different 86 yr old female donors with D-E of 3.3 and 6 hr with experimentation time of <24 hours and found that the 3.3 hr D-E interval globe exhibited superior viability (>70%), and nearly double the number of genes detected per cell (1301 vs 708 genes/cell). Therefore, globes with D-E times of < 6 hr and death to experimentation of <24 hr. Specifically, human globes from a 79 yr old female nonsmoker without AMD with a death to enucleation (D-E) time of 4.9 hr, a 67 yr old male smoker without AMD with a D-E of 3.8 hr, and a 67 yr old female smoker with early AMD (Minnesota Grade 2) with a D-E of 1.8 hr were immediately preserved on ice for overnight shipping to Hopkins, and dissected no later than 16 hr postmortem. After removing the anterior segment, the fundus was examined both with the retina and with the retina removed and classified using the Minnesota Grading Scale^105^. A 6 mm sample of the RPE/choroid in the macula was obtained using a 6 mm punch (Integra Lifesciences, Princeton, NJ) that was centered around the fovea. RPE/choroidal cells were dissociated using Papain Dissociation System (Worthington Biochemical, Lakewood, NJ) following the manufacturer’s instructions. Dissociated cells were resuspended in ice-cold PBS, 0.04% BSA and 0.5 U/μl of RNAse inhibitors. Cells were then filtered through a 50μm filter and processed for single-cell RNA-sequencing. Cell viability (>90%) was confirmed by negative staining with trypan blue.

### Nuclei Isolation Protocol

At 3, 6, and 10 days following the first injection, mice were sacrificed, and their eyes are enucleated. RPE/choroidal cells were scraped from the posterior eye cup and placed in cold 1ml 1X Dulbecco’s PBS. The nuclei from RPE/choroid (3 eyes from male and 3 eyes from female mice per condition) was prepared using the 10X Genomics “Nuclei Isolation from Complex Tissues for Single Cell Multiome ATAC + Gene Expression Sequencing” (10X Genomics, Inc., Pleasanton, CA) protocol. The RPE/choroid was centrifuged for 5 min with 500g at 4^0^C. The pellet was resuspended in 500ml of 0.1X Lysis buffer (10 mM Tris-HCl, pH 7.4,10 mM NaCl, 3 mM MgCl_2_, 0.1% Tween 20,0.1 % Nonidet P40,0.01 % Digitonin,1% BSA,1 mM DTT, RNase inhibitor 1 U/ml) and incubated for 5 min on ice, washed, and passed through a 70 μm Flowmi Cell Strainer followed by 40 μm Flowmi Cell Strainer, and centrifuged at 500 g for 5 min at 4^0^C. The resulting pellet was washed and resuspended in 0.1X Nuclei buffer containing trypan blue, and nuclei were counted with a Neubauer chamber. The obtained nuclei number were in the range of ∼2700 to 10000 nuclei/ml.

### SnRNA-seq and snATAC-seq library preparation and sequencing protocol

SnRNA-seq and snATAC-seq libraries were generated using the Chromium Next GEM single Multiome ATAC+Gene Expression kit protocol (10X Genomics, Inc.). Nuclei resuspended (10-15k) in 0.1X Nuclei Buffer were incubated with a transposition mix containing transposase. Gel Beads in Emulsion (GEMs) were generated by mixing transposed nuclei, barcoded Gel Beads, and partitioning oil on a chromium NEXT GEM Chip controller. After GEM generation, oligonucleotides included Illumina^®^ P5 sequence barcodes for ATAC library and cDNA included an Illumina^®^ TruSeq Read 1 (read 1 sequencing primer), 16 nt 10x Barcode (for GEX), 12 nt unique molecular identifier (UMI) and a 30 nt poly(dT) sequence. Magnetic beads (SILANE) were used to clean up barcoded products from the obtained GEM-RT reaction mixture. ATAC libraries were amplified with 10 PCR cycles and were sequenced on Illumina NovaSeq with 25,000 read pairs per nucleus. RNA libraries were amplified from cDNA with 14 PCR cycles and were sequenced on Illumina NovaSeq with 50,000 reads per nucleus.

### Bioinformatic analysis of snRNA-Seq and snATAC data

#### Mouse snRNA-seq

##### Quality control and removing cell doublets

Ambient RNA contamination were estimated and removed using R package DecountX^106^ based on RNA expression in the empty droplets. Low-quality cells were then removed based on following criteria: total RNA counts <1000 or >25000, total gene counts <200 or >10000, percentage of mitochondria RNA expression >50%. Cell doublets were then estimated and removed using R package DoubletFinder^107^. Only cells which passed the quality control were kept for downstream analyses.

##### snRNA-seq data integration, clustering, and cell type annotation

snRNA-seq samples were analyzed using R package Seurat v4.1.4^108^. All snRNA-seq samples were log normalized, scaled, and then integrated using Seurat functions (NormalizeData, ScaleData, FindIntegrationAnchors and IntegrateData). The integration was performed by using CCA to find integration anchors. The mouse 10-day control and 10-day CSC samples were used as reference for integration. The first 30 dimensions were used for further dimensional reduction and cell clustering. Cell clusters were then identified using “FindClusters” function with resolution of 0.24. Cells clusters were annotated manually based on the top 20 markers identified for each Seurat cluster.

To characterize RPE cells, we used the previously identified 154 RPE marker genes as reference^81^. Since the reference gene list was characterized for human RPE cells, we have modified the gene list by analyzing the expression of reference genes in 3 published mouse RPE cells^104,109,110^. The expression of 154 human RPE marker genes was compared between mouse RPE cells and other non-RPE cells. By applying a same cut-off (gene expression >0.5 in RPE cells and <0.5 in other cells) on the 3 datasets, we have identified 15 conserved RPE marker genes for mouse and used it for further RPE cells annotation (**Fig S1C**).

Differential expression analysis was tested between dedifferentiated RPE and RPE in control, CSC and aging mouse samples using Seurat “Findmarker” function with default parameters. Non-parametric Wilcoxon rank sum test was performed to identify differential expressed genes (DEGs) with a cut-off set as |avgLog2FC|>0.1 and false discovery rate (FDR) adjusted p value <0.05. After doublets and ambient RNA removal processes, we still found significant levels of rod and reticulocyte gene contamination in other cell types such as RPE and dedifferentiated RPE. Although RPE may also expressed low levels of rod genes such as *Rho*, these rod and reticulocytes genes were considered contamination because they were also highly expressed in the empty droplets. To avoid misinterpretation, we have identified the top 100 highest expressed genes in the empty droplets and manually removed them from the samples for downstream analysis.

The following gene lists were used to analyze EMT (Broad Institute’s GSEA EMT gene set, as previously published^75^), senescence (SenMayo^78^), and mitochondria (Broad Institute’s MitoCarta 3.0). For each cell, the expression of genes from a gene list (i.e.,EMT, senescence, and mitochondria gene list) were collected and summarized for each cell. The “EMT, senescence, and mitochondrial expression” of cells between dedifferentiated and healthy RPE was plotted. The final violin plot shows the distribution of “EMT, senescence, and mitochondrial expression” in cell clusters.

### Mouse snATAC-seq

All snATAC-seq samples were analyzed using Seurat v4.1.4^108^ and Signac v1.10.0^111^ packages follow the standard pipeline. ATAC peaks were called from each dataset separately using MACS2 peak calling function in Signac. Fragments from non-standard chromosome, Y chromosome and blacklist regions were removed from the peak-calling process. Called peaks from each sample were merged using “GenomicRanges: reduce” function to create a unified set of peaks. Counts of each peak in each sample was then quantified using “FeatureMatrix” function. Nucleolus with low quality peaks were excluded based on the following criteria: peak features >200 and <40000, peak counts >500 and <50000, nucleosome signal <2, Transcriptional start site (TSS) enrichment score >2, total number of fragments in peaks >200 and <60000. Since snRNA-seq and snATAC-seq data were generated from the same cell, each ATAC data was further filtered to keep nucleolus that also passed the snRNA-seq quality control. Frequency-inverse document frequency (TF-IDF) normalization was then performed to normalize sequencing depth differences across nucleolus using the “RunTFIDF” function in Signac. A singular value decomposition (SVD) was used for dimensional reduction on the TD-IDF matrix using all peaks (“FindTopFeatures” function, min.cutoff = “q0”). Since the first-dimension captures sequencing depth rather than biological variation, we used from the second to 30^th^ dimension to perform graph-based clustering, and non-linear dimension reduction for visualization using “RunUMAP”, “FindNeighbors”, and “FindClusters” function. As snRNA-seq and snATAC-seq shared the same cell barcode, snATAC-seq clusters were annotated using snRNA-seq annotations.

Differential accessible peaks (DAP) were analyzed using Seurat function “Findmarkers”. A logistic regression framework (test.use = “LR”) was used to determine differential accessible peaks with cut-off set as FDR-adjusted p value < 0.05. The DAPs were then annotated by its closest gene using Signac “ClosestFeature” function. If one DAP was annotated as a DEG and they changed the same direction, we considered the DEG was explained by chromatin changes.

KEGG enrichment analyses were done by using “kegga” and “goana” functions from R package edgeR^112^. A hypergeometric test was used to compare the number of DEGs in each KEGG term as compared to total number of genes in the term.

### Human scRNAseq

Single cell libraries were then prepared using the 10x Single Cell 3’ v2 Reagent Kits according to the manufacturer’s instructions and sequenced on an Illumina NextSeq500 using recommended sequencing parameters.

### RNA fluorescence *in situ* hybridization

After removing the neural retina, RPE flatmounts were fixed overnight at 4 °C in a 4% paraformaldehyde and incubated with RNAscope multiplex fluorescent reagent kit (Advanced Cell Diagnostics, Inc., Newark, CA) using the manufacturer’s protocol with mild modification. Briefly, the RPE flatmount was treated with H_2_O_2_ for 10 min and then protease IV solution for 30 min, both at room temperature, and then incubated for 2 hours at 40 °C with target probes (mouse *Ctsl* (Advanced Cell Diagnostics, Inc.) and *Ankfn1* (Advanced Cell Diagnostics, Inc.) in a HybEZ hybridization oven. RPE flatmounts were then treated with amplification reagent for 30 min at 40 °C and then incubated with RNAscope Multiplex FL v2 HRP-C1 at 40 °C for 15 min, TSA Vivid Fluorophore-520 at 40 °C for 60 min, and RNAscope multiplex v2 HRP blocker and then RNAscope Multiplex FL v2 HRP-C3, both for 15 min at 40 °C. The samples were then incubated with TSA Vivid Fluorophore 570, and with the RNAscope multiplex v2 HRP blocker both at 40 °C for 15 min. Finally, the RPE was stained with Anti ZO-1 antibody for 2 hr at 4 °C. RPE flatmounts were mounted on the glass slide with ProLong Gold mounting solution (Life Technologies, Inc., Carlsbad, CA). Images were obtained utilizing a confocal laser scanning microscope (LSM 880, Carl Zeiss Microscopy, Inc, White Plains, NY).

### TUNEL Assay

After RPE flatmounts underwent RNA *in situ* hybridization, followed by incubation with an anti-ZO1 antibody, apoptotic RPE cells were identified utilizing the Click-iT Plus TUNEL Assay kit supplemented with Fluor 488 dye (Invitrogen; Cat no. C10617) according to the manufacturer’s protocol.

## Funding

JTH (EY033765, EY031594, EY035805, RPB Stein Award, Robert Bond Welch Professorship), SW (EY035805, R01EY034571, BrightFocus grant in Age-related Macular Degeneration M2020166), DS (EY031594, Maryland Stem Cell Research Foundation Launch Grant, Foundation Fighting Blindness Free Family AMD Research Award, The Edward N. & Della L. Thome Memorial Foundation Awards Program in Age-Related Macular Degeneration Research, Frieda Derdeyn Bambas Endowed Chair), SB (R01EY036173), JQ (EY033765, EY031779, Karl Hagen Professorship); Core Grant, Wilmer Eye Institute, (EY001765); KS, YJ, MH, IP, TH, MC (none)

## Conflict of interest

JTH (SAB for Character Biosciences, Cirrus Pharmaceuticals, Seeing Medicines); SB (SAB and co-founder of CDI Labs). DS (Ikshana Therapeutics, Inc.)

LLMs were not used for this manuscript.

